# Contrasting climate signals between native and introduced annual and perennial floras

**DOI:** 10.64898/2026.07.31.741935

**Authors:** Elizabeth H. Wenk, William K. Cornwell, Ruby E. Stephens, David Coleman, Thomas Mesaglio, Isaac Towers, Sophie Yang, Daniel S. Falster

## Abstract

Annual versus perennial life histories represent a fundamental axis of plant strategy, with empirical evidence showing that the proportion of annuals is greater in hot, arid, or variable climates. Crucially, introduced species, a group that disproportionately includes annuals, are expanding in number and range across the globe. Whether introduced richness is governed by the same climatic controls as native richness remains untested at continental scales. For 19,299 native (12% annual) and 2,802 introduced (34% annual) Australian plant species, we show that native annual richness tracks climate far more tightly than introduced annual richness. Both floras largely follow the global pattern: the annual fraction rises in dry and seasonal climates. However, hotspots of native and introduced annual richness are completely different.

Introduced annuals (28% of all annual species) concentrate in wet, populated areas with low precipitation seasonality, whereas native annual richness is fairly evenly distributed, with the highest diversity in the seasonal tropics. The current continental-scale pattern is thus the sum of a long interaction between the native flora’s life histories and in situ climate, with dynamic, path-dependent introductions layered on top. As introductions and range expansions continue, they will reshape the annual-to-perennial balance along pathways of human influence.

## Introduction

Among the most fundamental distinctions in life history theory is whether plants complete their life cycle within a single year (annuals) or persist across multiple growing seasons (biennials and perennials)^1^. Theory predicts annual life cycles should predominate where environmental conditions limit the survival of roots and stems but not seeds, i.e., hot, arid, or climatically variable environments where persistence through dormant seeds is more advantageous than long-lived roots or stems^1,2^. Global analyses confirm this: annuals are disproportionately common in dry, hot, and variable climates^3,4^ and where favourable conditions are transitory^5,6^. These analyses have, however, treated plants as a homogenous group, not distinguishing between native species, whose distributions often reflect long-term climatic equilibrium, and introduced species, whose distributions are more dynamic and reflect human dispersal pathways.

There are several reasons why native and introduced floras would be distributed differently across the landscape. Native plants have mostly persisted through long-term climatic fluctuations in situ, allowing annual and perennial lineages time to adapt to local conditions or track suitable climates across the continent; their annual-to-perennial ratios should therefore closely reflect local climatic conditions in a manner consistent with the relative advantage of either strategy. By contrast, distributions of introduced species are co-shaped by the transport patterns of human movement^7–9^, combined with the niches inherited from their climate of origin^10^. Conflating the distributions of native versus introduced species risks misattributing the human-mediated dispersal signal to a universal annual climate response. It also masks if and how species introductions are dynamically shifting annual-to-perennial ratios across climate space.

Here we ask how annual prevalence shifts across climate space, and whether native and introduced species differ systematically in their climate-richness relationships. In doing so, we also extend recent global analyses relating life history to climate among herbs^4^ to include an entire continental flora. Australia’s flora offers a unique opportunity for such an analysis: it is a climatically diverse continent (Fig. 1a-c; Supplementary Fig. 1) with both a diverse native flora and a high number of introduced species (Fig. 1d) – and with datasets that include nativeness^11^ and life history^12^ scores for the entire flora. We predict that the native flora will reflect an equilibrium state that closely tracks climatic suitability, while the introduced flora richness will reflect human movement of plants and materials, plant dispersal, and climate in a dynamic fashion^13^. This leads to expectations that 1) the fraction of annuals in the flora will increase with aridity and precipitation seasonality, for both native and introduced floras; 2) the introduced flora richness will decline in dry and highly seasonal climates; 3) the fraction of the annual flora that is introduced will increase closer to population centres and in areas with more mesic conditions and less precipitation seasonality; and 4) the native flora richness will display a tighter climate coupling than the introduced flora. These predictions will allow us to assess the degree to which human-mediated dispersal is restructuring the annual-to-perennial balance on a continental scale.

**Figure 1.**
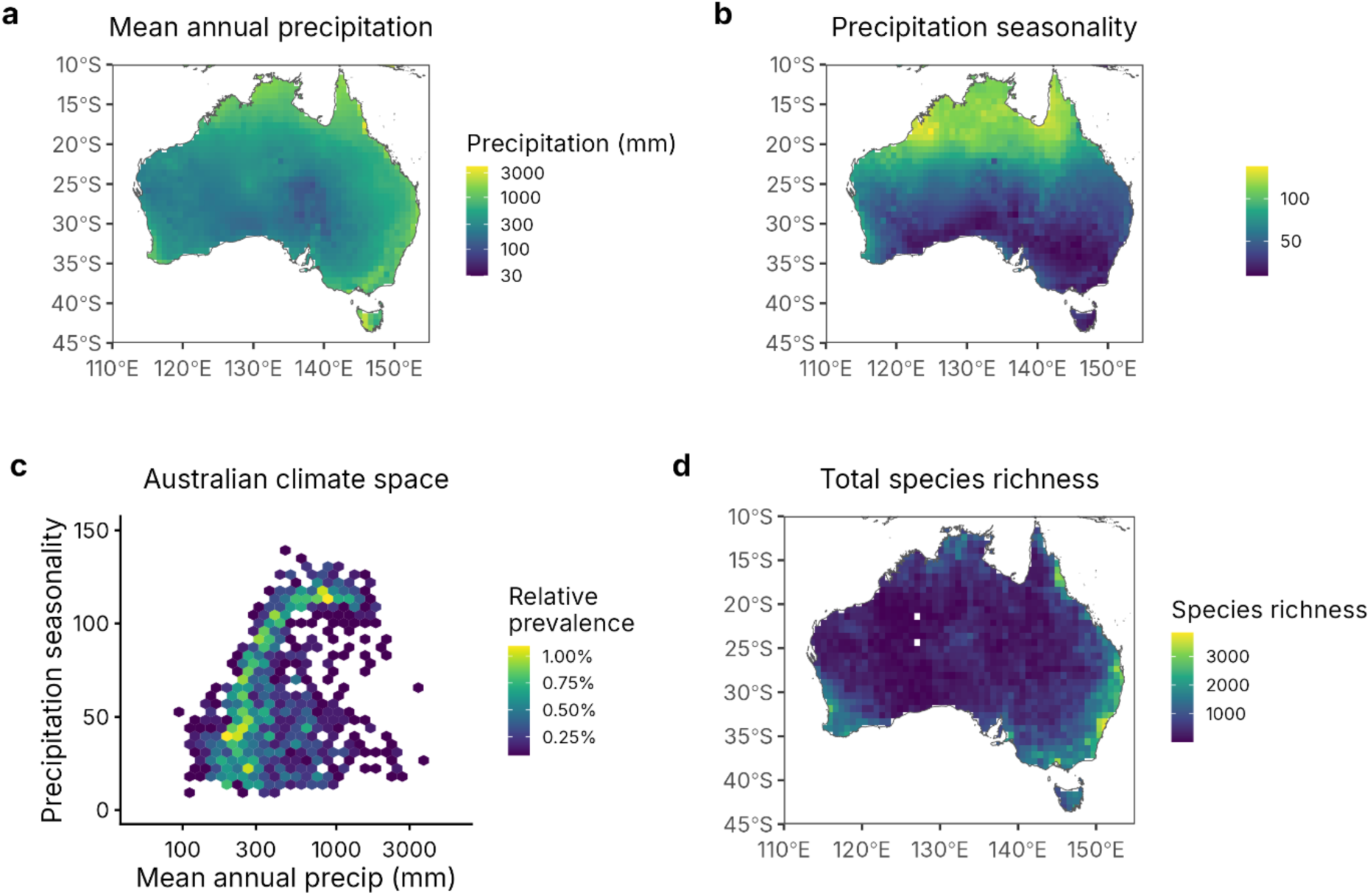
Gridded maps of Australia’s climate and plant richness including, a. mean annual precipitation (WorldClim variable bio12); b. precipitation seasonality (WorldClim variable bio15); c. prevalence of mean annual precipitation and precipitation seasonality values across Australia (see Supplementary Figure 1) for global climate space for these variables); d. total taxon richness (based on GBIF occurrence records). Gridded cells are 0.75° × 0.75°.

## Methods

### Dataset construction

We constructed a dataset of Australian vascular plant richness by integrating georeferenced occurrence records with taxonomic, life history, and climate datasets (Extended Data Fig. 1). We first assembled occurrence data for Australian vascular plants from the Global Biodiversity Information Facility^14^, starting from 24.4 million records and applying a sequential set of quality filters: restriction to Australian records from herbarium specimens and photographic vouchers, exclusion of records with coordinate precision poorer than 0.05° or coordinate uncertainty greater than 10,000 m, removal of records from before 1900 or with country-coordinate mismatches, and exclusion of two institutions (NHMUK, CJBG) known to have systemic georeferencing problems. We additionally applied CoordinateCleaner’s default validity checks to remove records with invalid, zero, or GBIF-headquarters coordinates. Records with missing coordinate-precision or coordinate-uncertainty values were retained. We augmented the package’s default reference tables with Australian capital cities and updated coordinates for one biodiversity institution that is incorrectly designated in CoordinateCleaner, CSIRO-Atherton, then removed records falling within 5 km of country/state centroids (22,373 records), 2 km of capital cities (25,656 records), and 2 km of biodiversity institutions (247,682 records). Records were retained at species rank and deduplicated by location × species × year, yielding 16.51 million cleaned occurrences for 23,626 species (Fig. 1e-f, Extended Data Fig. 1). Species names were aligned to the Australian Plant Census, Australia’s national consensus plant taxonomy reference, using the R package APCalign^15^; the APC includes consensus establishment status (native/introduced). Taxa that were indicated as being native to some parts of Australia and introduced elsewhere in the country were considered native for this study. The APC only includes taxa agreed by committee to be wild-growing in Australia. As such, using this resource likely undercounts introduced species because of the lag in the APC recognising new naturalisation events. Records for species not recognised by the APC were excluded (177,035 records). Life history status of each species (annual or perennial) was drawn from AusTraits^16^, explicitly using the life history scores in dataset “Wenk_2023”^12^, a 99% complete compilation of species’ life history values for the native and introduced flora of Australia, per the 2023-04-18 APCalign release. We chose to consider any plant that is capable of growing from seed and completing its reproductive life cycle within a year as possessing the capability to have an annual life history strategy. Therefore, species recorded in AusTraits as displaying both annual and perennial life history strategies were scored as annuals. Australia’s flora had 26,620 wild-occurring taxa at species rank (2026-03-26 APCalign release). Of these, 23,452 (88.1%) were native to at least somewhere in the country and 3,138 (11.9%) were introduced. Life history data were available for 25,588 of these species, with 21,922 (85.7%) species with no observations of an annual life history strategy and 3,666 (14.3%) species at least sometimes displaying an annual life history strategy (Fig. 2). While just 11.7% of native species are annuals, 32.9% of introduced species display an annual life history strategy.

**Figure 2.**
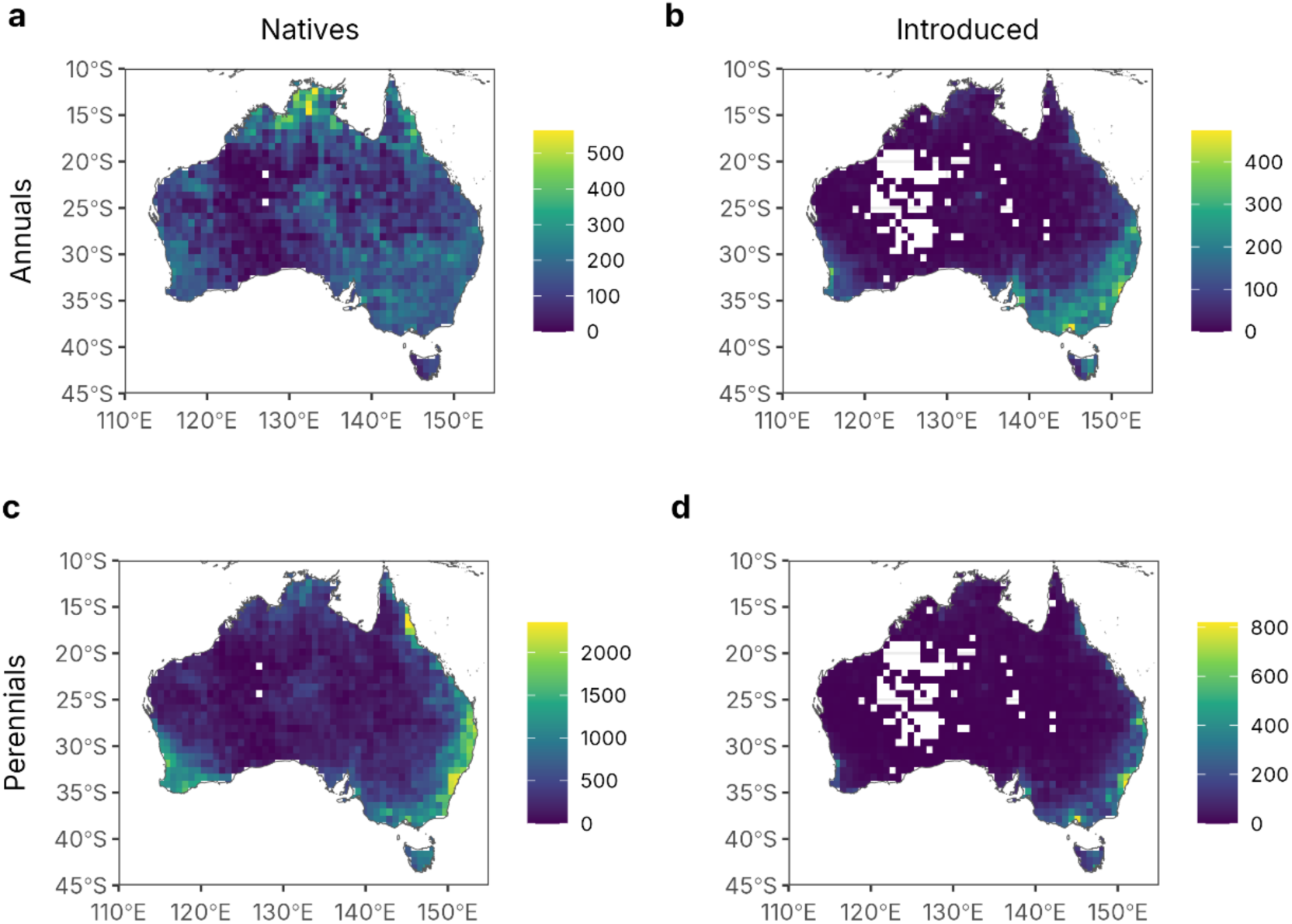
Gridded maps of species richness across Australia, specifically, a. native annuals; b. introduced annuals; c. native perennials; and d. introduced perennials. Gridded cells are 0.75° × 0.75°. Empty grid cells lack any observations for a given taxon grouping.

After intersecting the GBIF download with available life history and native status data, we retained records from GBIF for 22,101 of Australia’s 26,620 wild-occurring species, including 2,394 native annuals, 16,905 native perennials, 946 introduced annuals and 1,856 introduced perennials (Table 1).

**Table 1.**
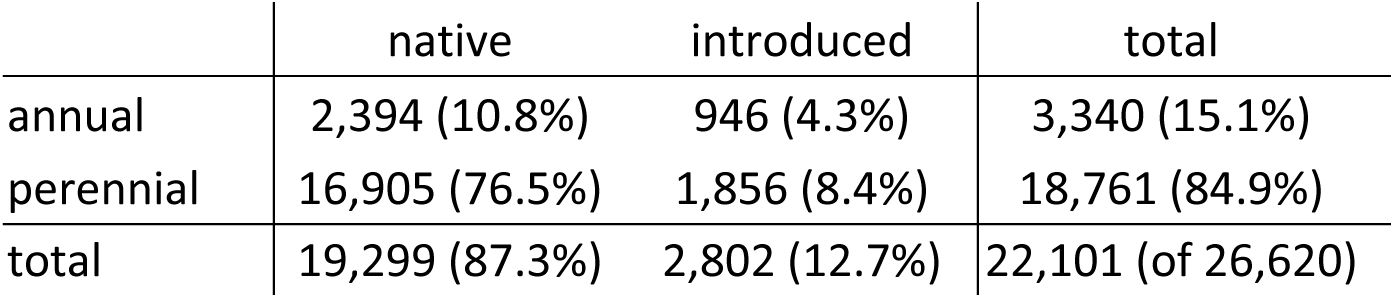
Percentage of sampled species that are annual versus perennial and native versus introduced. GBIF records and scores for both attributes were available for 22,101 of Australia’s 26,620 native and introduced plant taxa (83.0%). Percentages do not add perfectly across rows and columns due to rounding.

|  | native | introduced | total |
| --- | --- | --- | --- |
| annual | 2,394 (10.8%) | 946 (4.3%) | 3,340 (15.1%) |
| perennial | 16,905 (76.5%) | 1,856 (8.4%) | 18,761 (84.9%) |
| total | 19,299 (87.3%) | 2,802 (12.7%) | 22,101 (of 26,620) |

We sourced climate data from the WorldClim 2.1 bioclimatic layers at 2.5-minute resolution^17^, specifically using mean annual precipitation (bio12) and precipitation seasonality (bio15). We established a regular 0.75° latitude–longitude grid covering Australia (extent 112.5–154.0°E, 43.5–9.0°S; EPSG:4326) as our spatial analysis units and extracted the values for the climatic variables at the cell centroids (Extended Data Fig. 1). These values were linked to each occurrence record. Within each cell we tabulated the number of unique species in each of the four functional-origin groups (native annuals, native perennials, introduced annuals, introduced perennials), producing a cell-level richness dataset (Fig. 2). We also sourced national population data from the Australian Bureau of Statistics (www.abs.gov.au) and calculated the total population in each grid cell. To capture each cell’s exposure to human-mediated introduction, we derived a population accessibility layer based on grid cell population. For each 0.75° grid cell, accessibility was calculated as a Gaussian-kernel-weighted sum of square-root-transformed population density across surrounding cells, using a bandwidth of 250 km. Ocean cells were set to zero before smoothing so that coastal cells were not penalised for having an ocean neighbourhood, and the resulting values were applied to land only. Accessibility at each cell therefore reflects the combined, distance-weighted contribution of surrounding population centres as potential sources of species release and distribution.

### Analysis

To analyse how the fraction of annuals shifted with annual rainfall and rainfall seasonality we fit generalised linear models using the glmmTMB R package^18^. Log 10 mean annual precipitation (bio12), precipitation seasonality (bio15) and native status were included as predictors, using the model: fraction annuals ∼ log10(bio12)*native status + bio15*native status. The model was fit with a binomial response, weighted by total richness. Similar models were fit to analyse shifts in species richness across climate gradients, with species counts log10-transformed and fit with a Gaussian response. We filtered to cells with at least 500 occurrence records and at least two species in each life history class to restrict inference to well-sampled cells. Analyses with cells filtered to at least 100, 250 and 1000 occurrence records respectively are included in the Supplementary Material for comparison (Supplementary Figs. 2-4). Model predictions were generated by varying one climate variable across its observed range while holding the other at its sample mean.

To compare the goodness of fit for native versus introduced species against the climate variables, we fit four separate ordinary least squares regressions of log10 richness against log10(bio12) + bio15, one for each life history by species origin combination, and extracted the residual standard error from each. The residual standard error was used in preference to R², because R² is sensitive to the total variance in the response — groups with flatter responses yield lower R² even when the model fits equally tightly in absolute terms. The residual standard error therefore provides a directly comparable measure of typical absolute deviation between observed and fitted log10 richness across the four groups.

Sampling effort was heavily biased toward wetter, less seasonal locations, broadly tracking where human populations have historically concentrated (Supplementary Figs 5, 6). Because effort covaries with precipitation, observed richness confounds biology with detection probability; for non-native species, proximity to population centres additionally raises the likelihood of introduction. We therefore repeated the analysis on effort-standardised richness. All occurrence records in a cell were treated as a single pool, since a collector samples whatever is present regardless of origin or life history. For each of 20 random permutations of that pool we accumulated species separately for the four origin × life-history groups, fitted a power law (S = c · nᶻ) to each accumulation curve^19^, and evaluated it at a standardised effort of 500 records, taking the median across permutations as the point estimate. Cells with fewer than 500 records were dropped, so all values are interpolations rather than extrapolations. The three generalised linear models were then re-fitted to these effort-standardised values.

To test for an effect of proximity to population centres on the relative prevalence of introduced versus native species, we modelled the fraction of annual species that are introduced using a generalised linear model with a binomial error distribution, weighted by total annual species richness. The model included log10(mean annual precipitation), precipitation seasonality, population accessibility, and the interactions of population accessibility with each of the two climate variables as predictors: fraction introduced ∼ log10(bio12)* population accessibility + bio15* population accessibility.

The initial analysis pipeline was written entirely without use of AI. Subsequently, Claude (Anthropic) was used to help implement the kernel-smoothed population accessibility metric and rarefaction-based richness estimates, to enhance documentation of code, and to migrate analysis into a targets pipeline. All AI-assisted code was reviewed, tested, and validated by the authors before use.

## Results

### Geographic distribution of richness across life-history and origin groups

Native annuals, in contrast to both groups of introduced species and native perennials, displayed a mostly quite even distribution across the continent, but with high richness in the Top End of the Northern Territory (Fig. 2a). Introduced annuals had the highest richness near population centres, and relatively high richness throughout the southeast part of the continent and Tasmania more broadly (Fig. 2c). Both native and introduced perennials had the highest richness close to the coast, with native richness strongly aligned with known diversity hotspots such as tropical Far North Queensland and southwestern Western Australia (Fig. 2b), while introduced perennials were more aligned with coastal population centres and their richness declined rapidly inland (Fig. 2d).

### The fraction of annuals declines with precipitation and varies with precipitation seasonality

Climate data layered over these geographic patterns showed how the fraction of annuals in the native versus introduced flora shifted in response to log10 mean annual precipitation (WorldClim variable bio12) and precipitation seasonality (WorldClim variable bio15) (Fig. 3, Fig. 4, Supplementary Table 1). The fraction of annuals was lower in wetter locations (z=-67.04; p < 0.001), lower among native species (z=-29.31; p < 0.001), and not significantly correlated with precipitation seasonality (z=1.30; p=0.195), although significant interaction terms indicated different responses between native and introduced species to climate variables (Supplementary Table 1a). The fraction of annuals in both floras increased as precipitation decreased, although the fraction of annuals was much higher in the introduced flora; in addition, the response to mean annual precipitation was lower for native than for introduced species (Fig. 3a-b, Fig. 4a-d, Supplementary Table 1a). For precipitation seasonality there was a non-significant overall effect, but strongly significant interaction; the fraction of annuals in the native flora increased with increasing precipitation seasonality, while for the introduced flora there was no shift in the fraction of annuals (Fig. 3a-b, Fig. 4a-d, Supplementary Table 1a).

**Figure 3.**
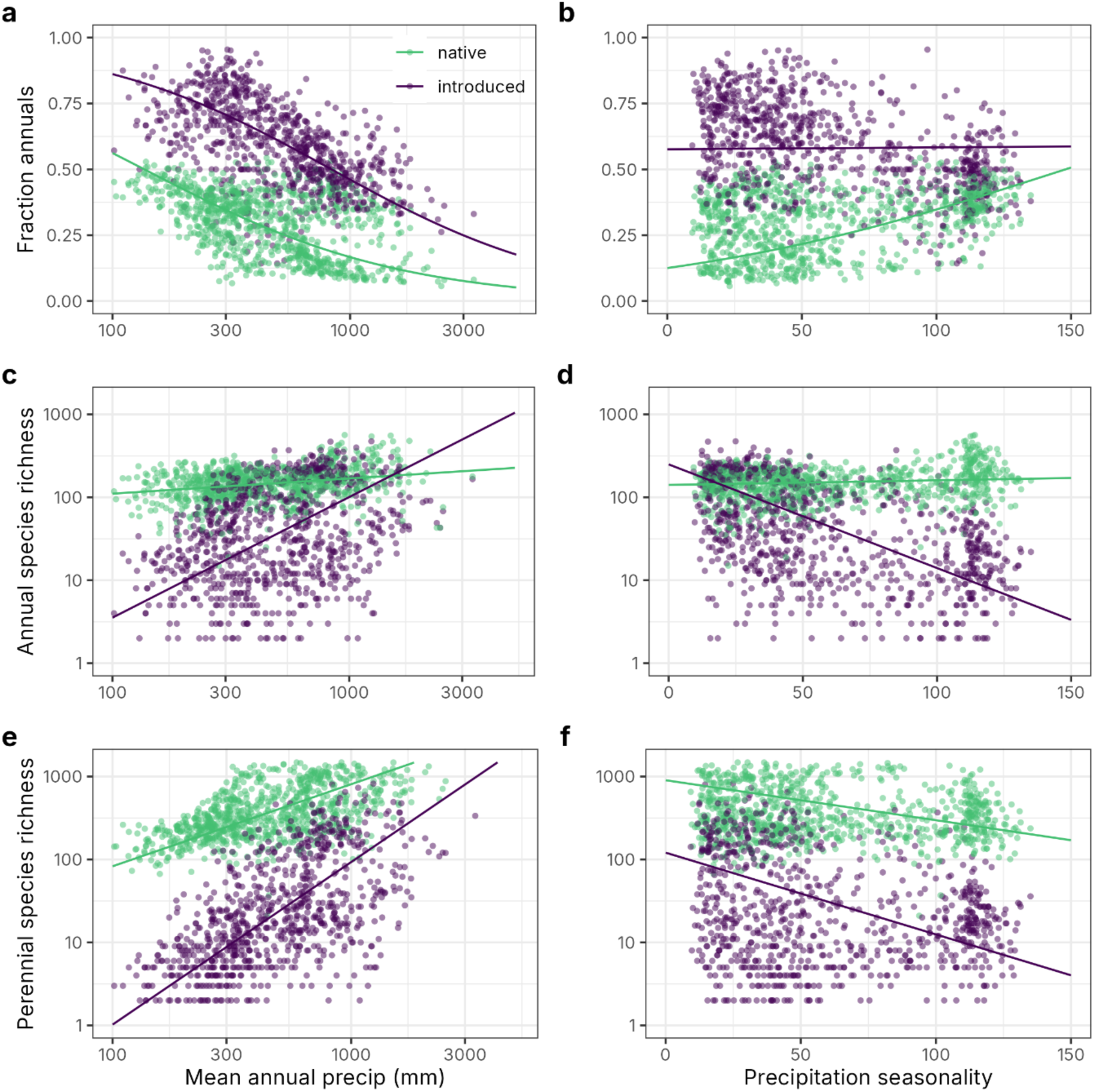
Native versus introduced species richness in each grid cell (0.75° × 0.75°) by climate, specifically: a. fraction of annuals with mean annual precipitation; b. fraction of annuals with precipitation seasonality; c. annual richness with mean annual precipitation; d. annual richness with precipitation seasonality; e. perennial richness with mean annual precipitation; and f. perennial richness with precipitation seasonality. Grid cells with fewer than 500 records were omitted.

**Figure 4.**
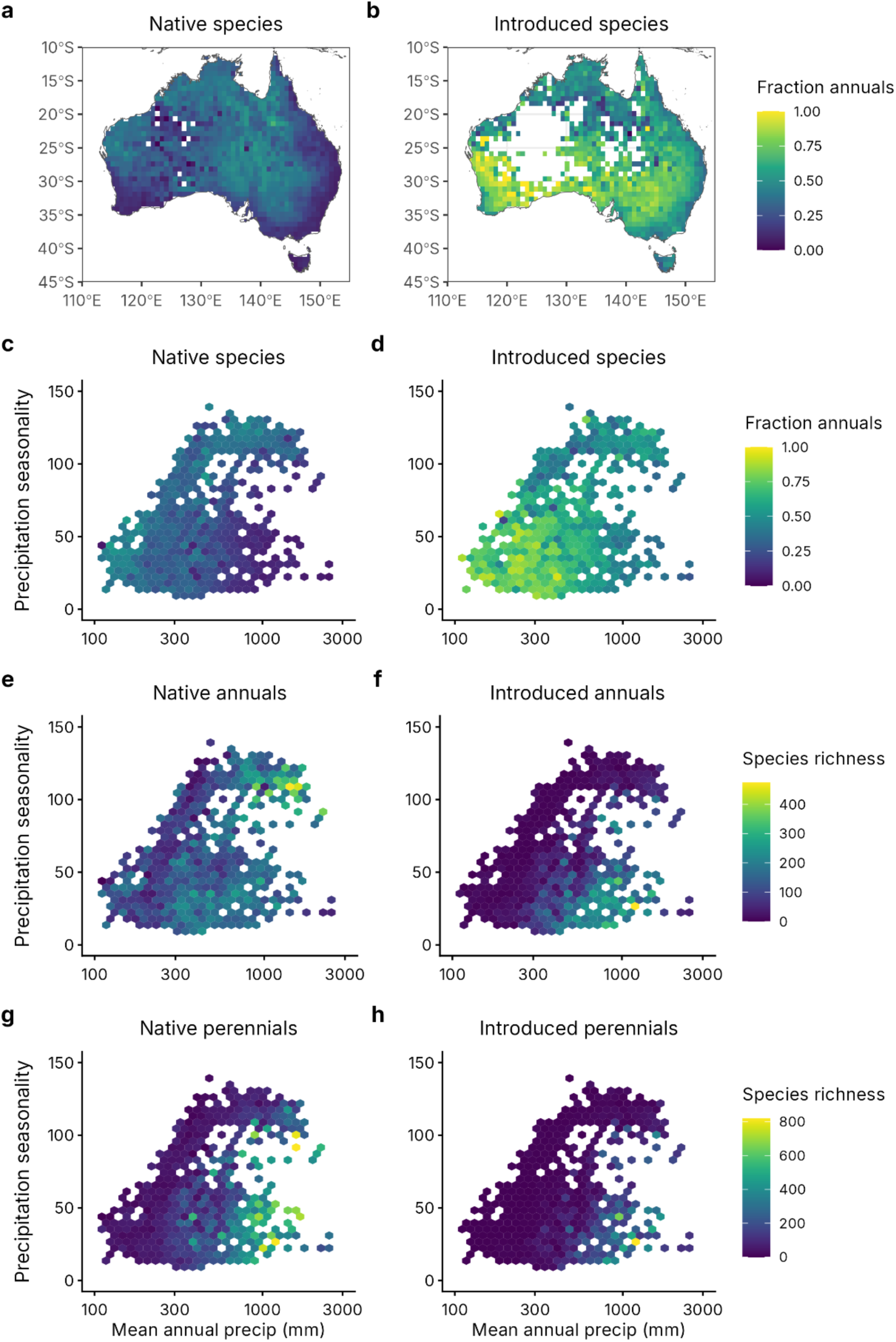
Hexbin plots and gridded maps showing fraction annuals and species richness across Australia’s climate space. Specifically, a. fraction native species that are annuals across Australia; b. fraction introduced species that are annuals across Australia; c. fraction native species that are annuals across Australian climate space; d. fraction introduced species that are annuals across Australian climate space; e. native annual richness across Australian climate space; f. introduced annual richness across Australian climate space; g. native perennial richness across Australian climate space; and h. introduced perennial richness across Australian climate space. Grid cells are 0.75° × 0.75°. Empty grid cells lack any observations for a given taxon grouping.

### Richness components underlying the annual fraction

Because the fraction of annuals is the ratio of annual to total richness, its response to climate is governed by the difference in slopes between annual and perennial richness rather than by either alone. The richness of annuals and perennials in the native and introduced floras showed different responses to mean annual precipitation and precipitation seasonality. For the pair of models fit to annual and perennial richness, we found strong evidence for all fixed effects and their interactions (Fig. 3c-f, Supplementary Table 1b-c). Native annual richness increased only slightly across the continental mean annual precipitation gradient, while native perennial richness was higher in wetter locations, leading to the observed decline in native annual fraction in wetter regions (Fig. 3a, c, e, Fig. 4 e, g, Supplementary Table 1). Meanwhile, for the introduced flora, the richness of both annual and perennial species increased strongly with precipitation although perennial species richness showed a steeper response, leading to the observed decline in annual fraction with precipitation in the introduced flora (Fig. 3a, c, e, Fig. 4 f, h, Supplementary Table 1).

For precipitation seasonality, native annual richness did not shift significantly across the gradient, while native perennial richness declined slightly, leading to the steep increase in annual fraction with increasing precipitation seasonality. Introduced annuals and perennials showed similarly steep declines in richness as precipitation seasonality increased leading to the lack of a shift in introduced annual fraction across this climate gradient (Fig. 3b, d, f, Supplementary Table 1).

### Native climate–richness relationships are tighter than introduced

Native species richness was more tightly predicted by climate than introduced richness: RSE was lower for native than introduced annuals (0.187 vs. 0.354) and perennials (0.215 vs. 0.376, see also Fig. 3).

### Human accessibility shapes the introduced fraction of annuals

Among Australia’s annual species (native and introduced combined), the introduced fraction increased with higher annual precipitation (z = 61.95; p < 0.001), lower precipitation seasonality (z = −74.42; p < 0.001), and greater population accessibility (z = 42.32; p < 0.001) (Fig. 5, Supplementary Table 2). There was a significant negative interaction between annual precipitation and population accessibility (z = -41.24; p < 0.001); the positive effect of high annual precipitation on the fraction of introduced annuals was accentuated in areas with low population accessibility. Meanwhile, there was a significant positive interaction between precipitation seasonality and population accessibility (z = 46.58; p < 0.001), indicating that the impact of precipitation seasonality on the fraction of introduced annuals was highest in areas with low population accessibility. Overall, in climatically less favourable locations, whether those with lower precipitation or higher precipitation seasonality, the effect of proximity to population centres was accentuated (Extended Data Figs. 3-4).

**Figure 5.**
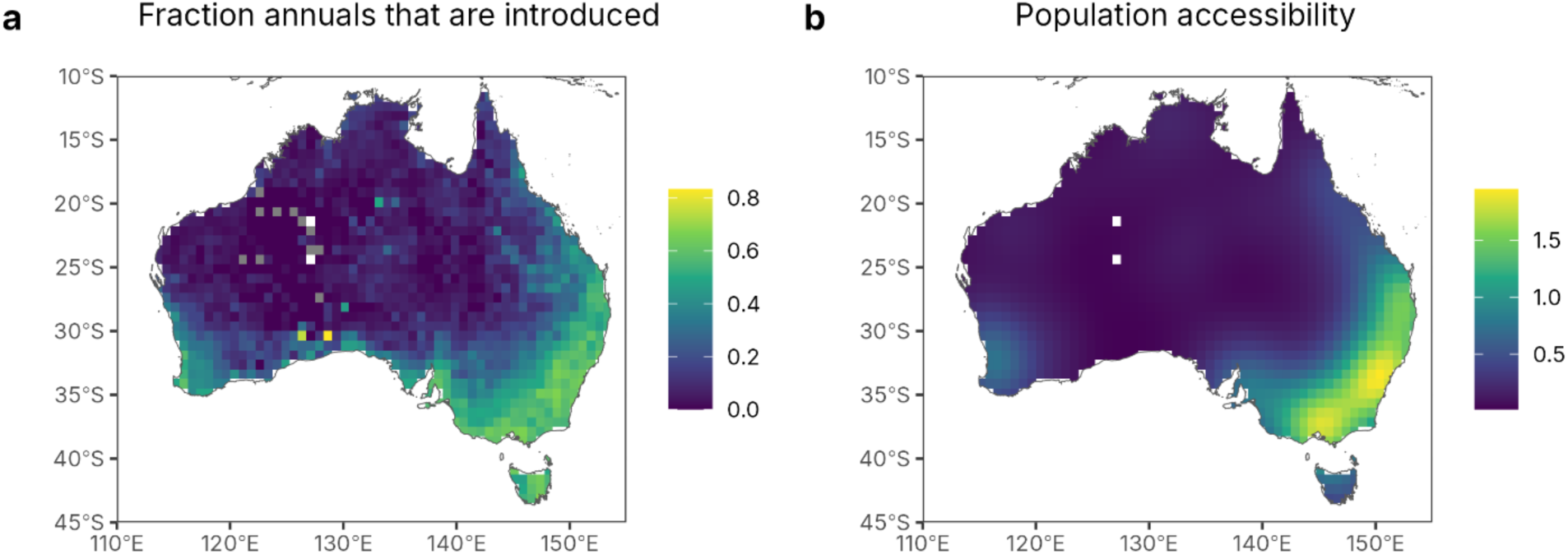
Gridded maps comparing a. fraction of annual species that are introduced, and b. population accessibility. Grid cells are 0.75° × 0.75°.

### Robustness to sampling effort and rarefaction

Because overall sampling effort is highest in high-precipitation, low-seasonality locations (Supplementary Fig. 5), we repeated these analyses using effort-standardised species counts. The patterns in the fraction of annuals across climate gradients held qualitatively in response to this correction (Extended Data Fig. 2a–b, Supplementary Table 3). Standardising to a common effort of 500 records reduced richness in every species grouping but did so unevenly along both gradients: the median value for rarefied richness across species groupings was 13–18% of the raw count in the wettest quartile of cells but 44–56% in the driest, and 21–24% in the least seasonal quartile but 35–44% in the most (Extended Data Fig. 2c–f). The resulting shifts in slope were consistent across all species groups and both climate variables: every relationship with mean annual precipitation shifted downward, by 0.00066–0.00078, and every relationship with seasonality shifted upward, by 0.0053–0.0068 (Supplementary Table 4). Rarefaction therefore preserved the differential responses across life-history and origin groups, while showing that the apparent richness advantage of wetter, less seasonal cells is substantially a product of their greater sampling effort. Two of the eight relationships changed sign under the correction — native annual richness against precipitation, and native perennial richness against seasonality — but both involve slopes indistinguishable from zero in the raw models.

To avoid drawing inferences from under-sampled cells, the analyses above were restricted to grid cells with more than 500 occurrence records (904 cells). We repeated the tests at three additional cut-offs (100, 250 and 1,000 records; Supplementary Figs 2–4). Including low-effort cells worsened model fit, with the residual standard error for annual richness rising from 0.26 at the 1,000-record cut-off to 0.32 at 100. Seven of the eight climate–richness relationships kept both their sign and their approximate magnitude across the full range of cut-offs. The exception was native annual richness, whose weak relationships with both climate variables proved cut-off sensitive; its slope against seasonality changed sign between the 250- and 500-record cut-offs. Meanwhile, the 1,000-record cut-off gave results close to the 500-record cut-off, with 747 rather than 904 cells.

## Discussion

For native species, continental data on Australian plants support both theory^1,2^ and recent empirical work (focused on herbs)^4^ showing that lower or more variable precipitation promotes the annual habit, particularly as a fraction of the total flora. Such climate-associated patterns in life history are often taken as a guide for how biodiversity will reorganise under climate change. Yet we find a contrasting diversity signature in introduced species, suggesting that the reconfiguration of the flora is not following these predicted climatic patterns. Specifically, the introduced annual fraction is higher in more mesic, less seasonal areas and closer to population centres — presumably reflecting the combined signatures of climate-of-origin, species arrival, and landscape disturbance. Together, these results suggest that future biodiversity will be shaped as much by human movement of plants across continents as by climate shifts.

The maintenance of native annual richness across climate gradients confirms that, although the annual habit may be most advantageous in dry and unpredictable environments (relative to the perennial strategy), it is nonetheless ecologically successful across Australia’s entire climatic space. Native annuals encompass a remarkable diversity of trait constellations, each conferring fitness under different conditions (Extended Data Fig. 5). Globally, traits that raise fitness in arid and unpredictable environments include environmentally triggered germination cues, long-lived soil seedbanks^20,21^, bet-hedging through plasticity in seed germination^22–25^, high dispersal^26^, and timing reproduction so the growing season remains favourable until seed set^27–29^. Australia’s native annuals span this breadth: edaphic specialists confined to ephemeral pools, salt lakes, or unusual soils; fire ephemerals whose germination is tightly linked to post-fire conditions^20,21,30^; but there are also disturbance specialists characterised by small seeds, high seed output, cheap tissues, determinate growth, and rapid transition to reproduction. This diversity of strategies lets native annuals occupy niches across the continent’s full climate space — as reflected in their consistent richness along climate gradients (Fig. 3a, c, Fig. 4).

Introduced annuals show the opposite pattern. Where native annuals occupy a wide variety of narrow, specialised niches across macroclimate gradients, introduced species are, broadly speaking, generalists of human landscapes, tracking both the benign macroclimates people favour and the ruderal opportunities their disturbance creates. Accordingly, the richness of introduced species, annual and perennial alike, declines steeply with aridity, precipitation seasonality, and distance from population centres, and is consistently low in arid, highly seasonal locations (Figs. 2b, d, 3d, f, Fig. 4). There were 157 grid cells (0.75° × 0.75°) in which native species but no introduced species were recorded — all in climatically unfavourable, sparsely populated locations. This makes sense: Baker’s^31^ “ideal weed” is defined by ruderal traits that aid establishment — small seeds, rapid reproduction, high seed output^32^ — not by adaptations to recurring drought or unpredictable rainfall. Also, introduced distributions correlate with known points of first record^33,34^, and most problematic invaders arrived through the horticultural trade^13^. Moreover, introduced species tend to establish where climate matches their traits and native range^10,35^, mesic, relatively aseasonal climates best suit the low-dormancy, fast-growing, ruderal traits typical of weeds. Such species may establish briefly in harsher locations, but decline or vanish after dry years, since they are more impacted by drought compared to co-occurring natives^36^ and are less likely to bet-hedge^37^. Specialists of rare environments, such as edaphic endemics^38^ or post-fire ephemerals^30^, rarely move across continents, probably because human dispersal seldom delivers propagules to the precise soils or disturbance regimes they need.

Climate shapes the richness of both native and introduced species (Fig. 2), but that shared signal conceals a macroecological mismatch, revealed by how tightly each group tracks climate. Native richness is far more closely coupled to climate than introduced richness: residual standard errors are roughly half as large for natives. The weaker climatic signal for introduced species points to a considerable degree of disequilibrium — their distributions have yet to fill, or settle into, the climate space their traits would predict. Not only are introduced species largely absent from climatically unfavourable environments, but they also occur less predictably across the full range of climate space. In locations close to population centres with favourable climate, introduced annual richness approaches that of natives (Fig. 3, Extended Data Figs. 3, 4), while in climatically similar but less accessible locations, richness declines sharply. The weak coupling could reflect introduction history alone, in which case it should erode as species spread from their introduction points; or it could represent a real match between the ruderal traits of introduced annuals and the disturbance regimes of densely populated landscapes, in which case introduced annual richness will continue to track human population density.

Viewing richness shifts through the distribution of annuals alone masks a key pattern: perennials respond just as strongly to climate, and introduced perennials show by far the steepest response of any group (Fig. 3e,f). In arid or variable environments, a high annual fraction often reflects declining perennial richness rather than rising annual richness: woody perennials lose viable trait combinations as conditions harshen^39,40^, while annuals are less affected^28,41,42^. Presumably, this is because the environmental conditions that annual plants experience in their vegetative phase can differ markedly from those of perennials even in the same location; germination cues paired with a short lifespan ensure that annuals experience a comparatively wetter climate during their life cycle, even in dry areas. Recruitment in long-lived perennials compounds this: establishment is episodic, succeeding only in rare events across decades. Both native and introduced perennials became richer under more favourable precipitation, but the introduced response was about twice as steep (Supplementary Table 1; Figure 3e, f). This fits the idea that introduced perennials are the group least able to establish in arid, highly seasonal locations: there, the rare recruitment windows are complex and infrequent.

The climate–fractional annual contrasts (Fig. 3a-b) are generally robust to sampling bias. However, consistent undersampling in climatically unfavourable locations hints at undetected species richness (Supplementary Figures 5, 6). Rarefaction analysis confirmed that the direction of the fraction-of-annuals responses were unchanged for both native and introduced species across both climate variables, while richness slopes flattened approximately equally across all four groups (Extended Data Fig. 2, Supplementary Tables 3, 4), as rarefaction analyses led to a greater reduction in richness in favourable versus unfavourable environments. That symmetry matters: whatever species remain undetected in Australia’s less-sampled arid and seasonal environments, they are distributed proportionally across native and introduced, and annual and perennial groups. The implication is twofold — native diversity in unfavourable climates is almost certainly higher than recorded, but so too may be introduced richness, suggesting that introduced species may be filling climate space more than current occurrence data reveal. The rarefaction method cannot predict actual richness anywhere, since capping richness in well-sampled locations inherently flattens slopes; it underscores the importance of continued targeted survey effort in remote environments.

Global analyses of life-history biogeography have largely ignored species origin, an omission that distorts our understanding of how annuals are distributed across climate space. Poppenwimer *et al.*^4^, for instance, recorded neither native nor introduced status and excluded woody plants entirely. A continental-scale analysis allows for this distinction, and we show here that some patterns hold across origin groups: the fractional response of annuals to climate was broadly similar for native and introduced species in both their analysis and ours. But for a key driver, richness, species origin is a confounder. Our results do not support the inference that human disturbance broadly favours annuals; it favours the establishment of introduced annual species. Native annual diversity is as high in the most invaded locations as the most pristine, indicating that introduced species likely establish through propagule pressure and climate matching rather than by filling vacant niche space^8,43^. Conflating native and introduced responses risks predictions of future annual spread that do not apply equally across species and places. Australia is among the highest recipients of introduced plant species globally, making it an especially instructive system for understanding how introduced species reshape the annual-to-perennial balance.

Species richness tells us how many introduced taxa have established, but richness is not abundance — and abundance drives ecological impact. A single introduced species can transform an ecosystem if it reaches high local abundance. The clearest cases come from climatically unfavourable environments with low introduced species richness, such as buffel grass (*Cenchrus ciliaris*) and tamarisk (*Tamarix aphylla*) in Australia’s arid zone, and cheatgrass (*Bromus tectorum*) in the western United States^8,44,45^. While the relationship between richness and abundance is complex and beyond the scope of this paper, the scarcity of introduced taxa in harsh climates should not be mistaken for ecological safety^46^. Looking forward, forecasting the future distribution of annual versus perennial life histories will require integrating three fields of knowledge: life-history theory^2^, particularly the seed-to-rhizome survival ratio across climates and habitats; regeneration ecology, including the episodic, disturbance-driven nature of perennial recruitment; and the ecology of introduced species.

Australia’s annual-to-perennial balance is the sum of two very different histories: a native flora whose life histories have equilibrated with climate over millions of years, and an introduced flora assembled over two centuries along pathways of human movement. Whether the two will ever converge is unresolved. Many introduced taxa struggle to persist in conditions where bet-hedging pays off, conditions predicted to become more prevalent including fire, drought, and floods^47,48^. Our analysis is of Australia, and a continental-scale lens is what makes near-complete data on both species’ origin and life history attainable in the first place, but the principles can be generalised across the globe. The annual-to-perennial balance on any continent plays out on a climatic stage, with species as the actors moving across it as it changes — the “conserving nature’s stage” framing of Lawler *et al*.^49^. What that framing assumes to be slow, however, is now fast: humans are not just shifting where the existing actors stand but adding new ones to the cast, introducing plants far more quickly than the stage itself is changing. That is what makes the annual-to-perennial balance one of the most dynamic aspects of global plant biodiversity in the coming century and introduced species its least predictable driver.

## Acknowledgements

We thank members of the Cornster Lab at UNSW for valuable discussions and feedback on an early draft. The authors have no conflicts of interest to declare.

## Data Accessibility and Reproducibility

The code to reproduce the results can be found at https://github.com/traitecoevo/cornsters_australian_annuals. This paper uses species occurrence data from GBIF (https://doi.org/10.15468/dl.dhnekr) and trait data from the AusTraits database (Falster et al. 2021, https://doi.org/10.5281/zenodo.7583087).

## Author contribution statement

Will Cornwell, Daniel Falster, Isaac Towers and Elizabeth Wenk conceived the idea for the paper and did the analyses. Elizabeth Wenk and Will Cornwell wrote the initial draft. All authors contributed to figures and tables for the manuscript. All authors critically reviewed and edited drafts of the manuscript. All authors reviewed the final paper for publication.

## Extended Data

**Extended Data Fig. 1.**
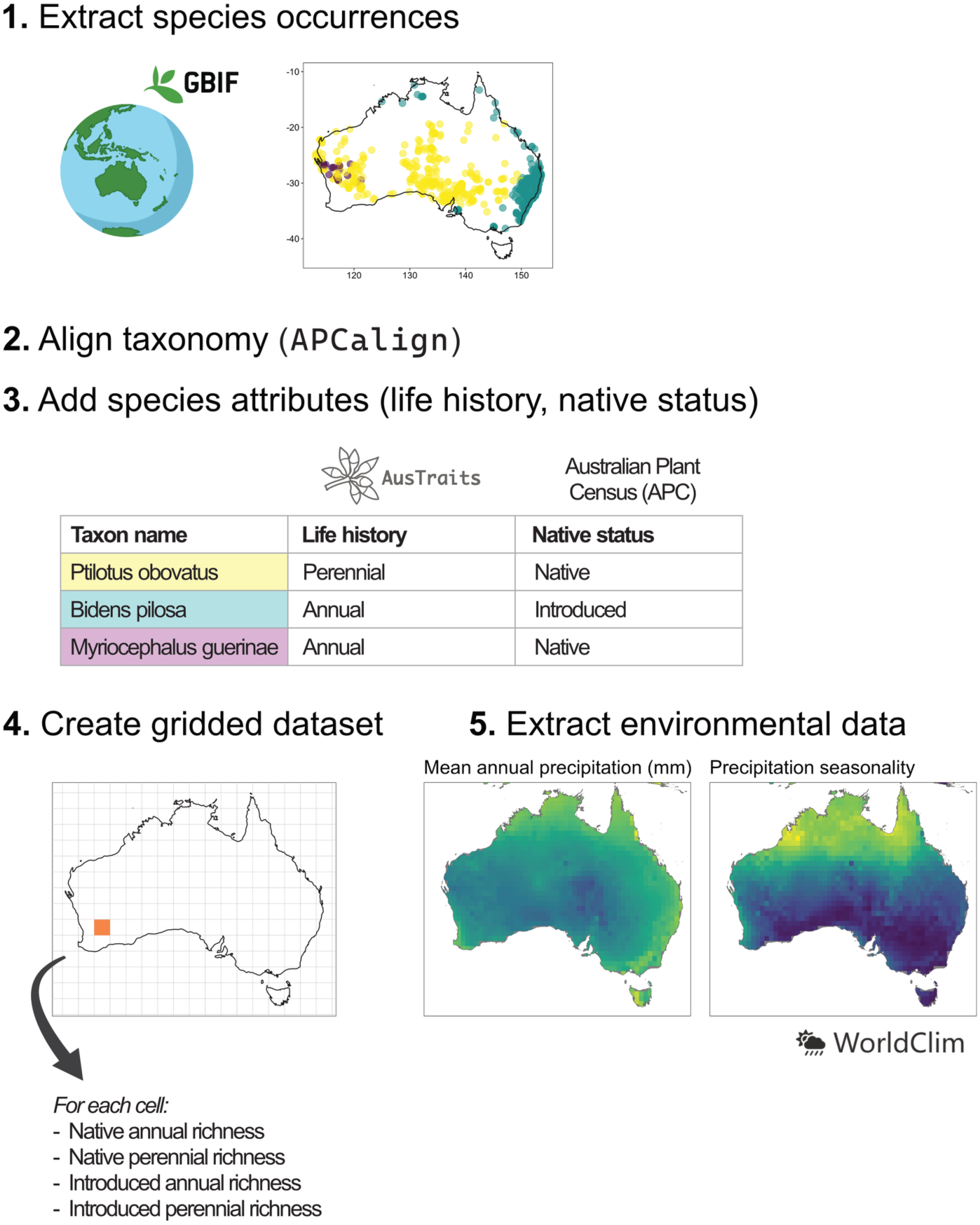
Workflow for synthesising occurrences with trait data and climate data. Step 1. Extract occurrence records from GBIF; Step 2. Align to APC taxonomy using APCalign; Step 3. Extract life history data from the AusTraits database and native status from APCalign; Step 4. Produce a gridded map of the number and fraction of native and introduced annual and perennial species across Australia; and Step 5. Extract climate data from WorldClim and join to gridded occurrence data.

**Extended Data Fig. 2.**
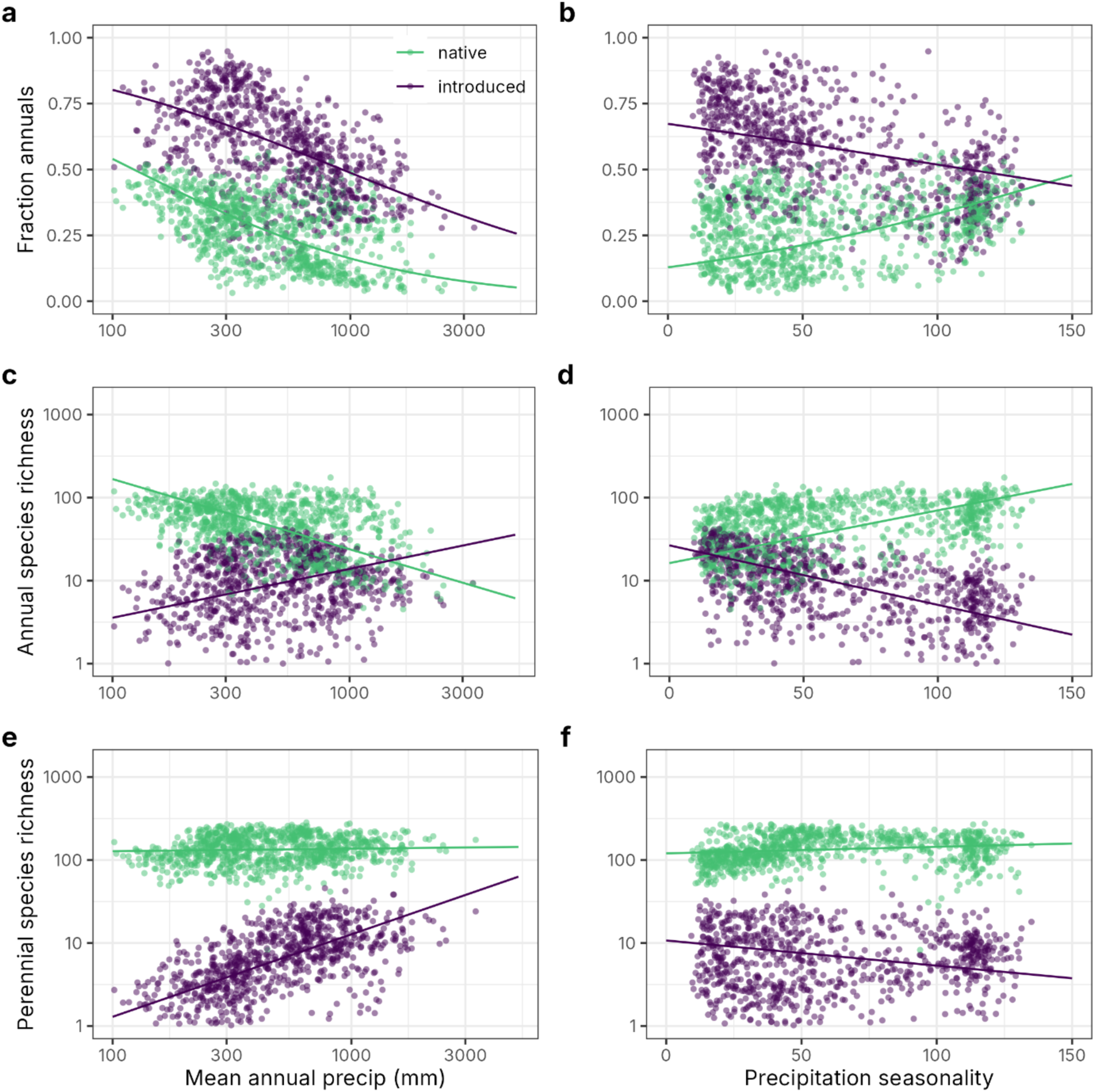
Figure 3 repeated, using rarefaction. Grid cells with fewer than 500 records were omitted.

**Extended Data Fig. 3.**
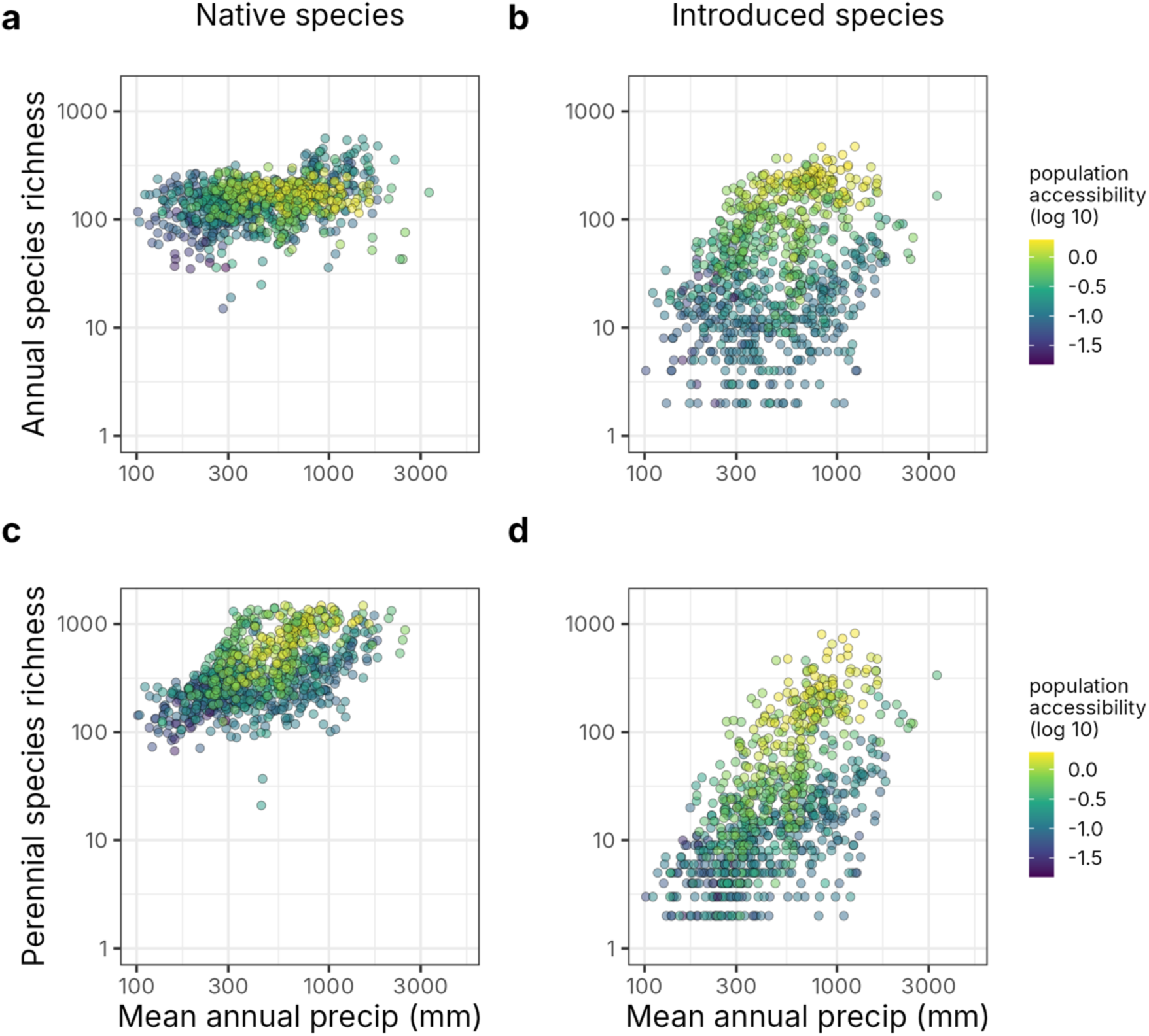
Shifts in species richness across the mean annual precipitation gradient, with values shaded by population accessibility.

**Extended Data Fig. 4.**
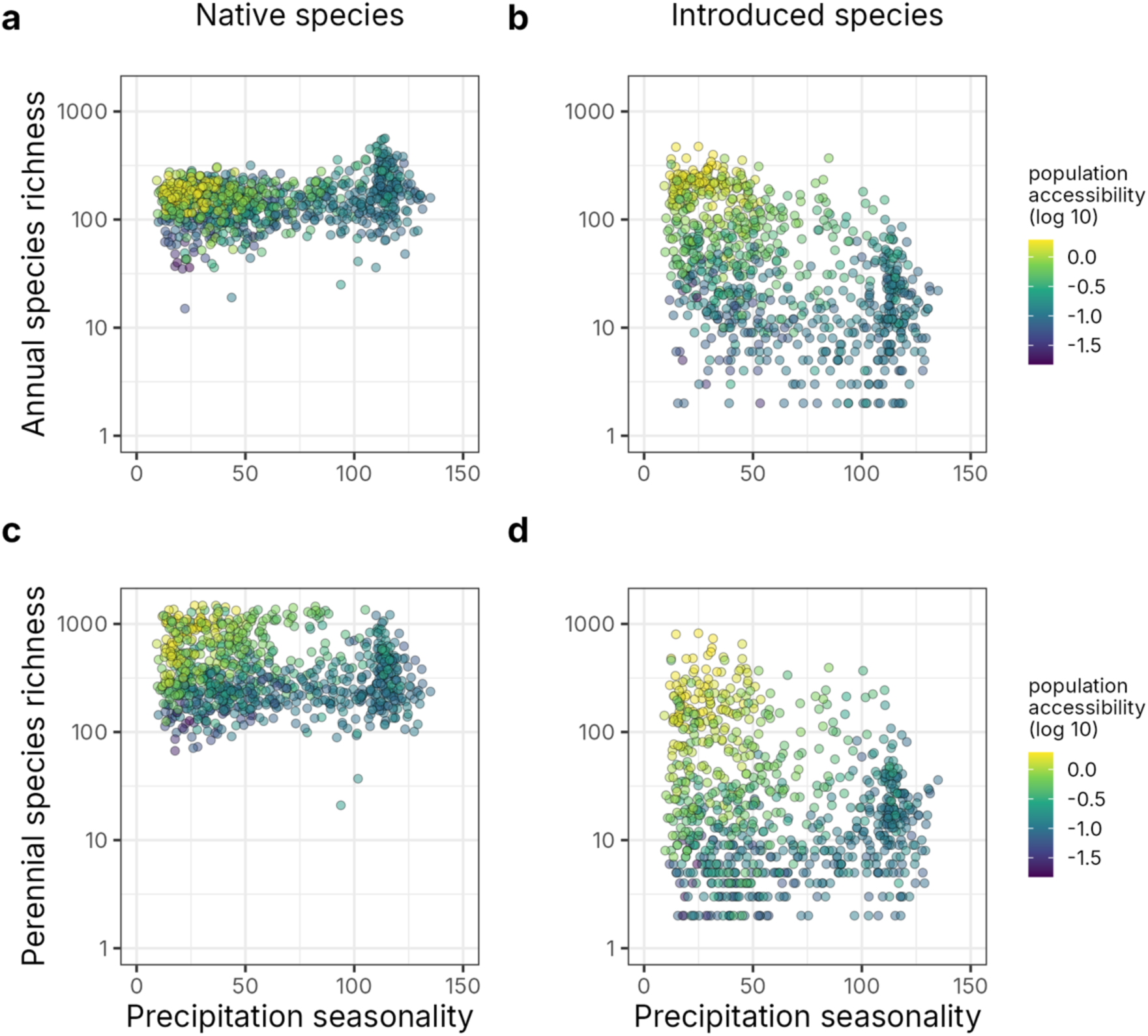
Shifts in species richness across the precipitation seasonality gradient, with values shaded by population accessibility.

**Extended Data Fig. 5.**
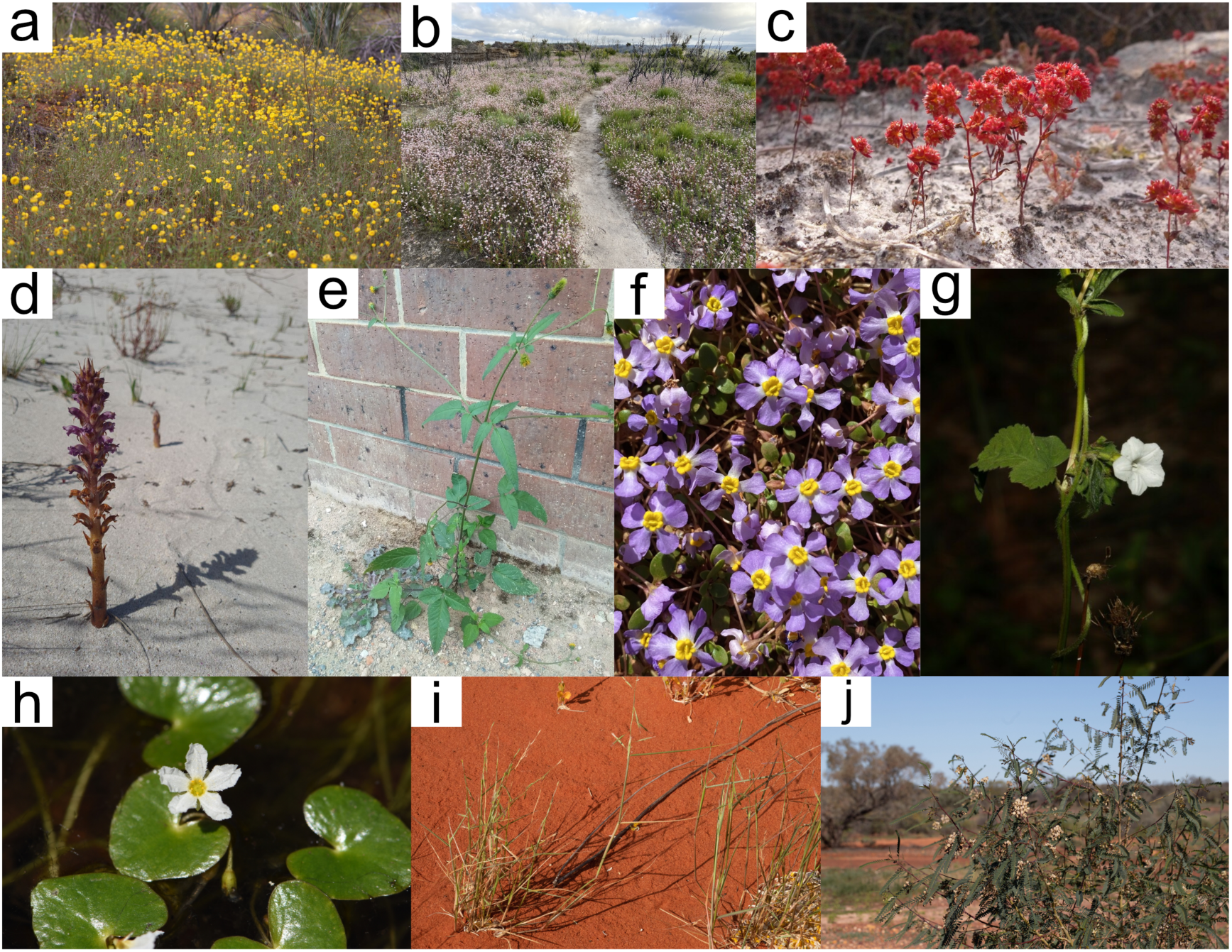
Illustrating the diversity of annual plant strategies and forms found in Australia. a. Rare paper daisy native to arid Western Australia, Myriocephalus guerinae (©Arthur Chapman); b. Fire ephemeral, Actinotis forsythii (©Margaret Sky); c. Introduced South African succulent, Crassula glomerata (©Bo Janmaat); d. Parasitic annual, Orobanche cernua (©Anton Hooft); e. Urban non-native annual, Bidens pilosa (©’insiderelic’); f. Undescribed/phrase name annual, Peplidium sp. C Evol. Fl. Fauna Arid Aust. (N.T. Burbidge & A. Kanis 8158) (©Loxley Fedec); g. Climbing annual, Ipomoea plebeia (©Greg Tasney); h. Aquatic annual, Nymphoides minima (©Alistair Smith); i. Annual grass, Paspalidium reflexum (©Euan Moore); j. Rare example of an annual shrub, Sesbania benthamiana (©Kym Nicolson). All photographs from iNaturalist.

## Supplementary Figures & Tables

**Supplementary Table 1.** Responses of a. fraction annuals; b. annual richness; and c. perennial richness to log10(bio12) (mean annual precipitation, mm), bio15 (annual precipitation seasonality), and native status (native, introduced). The model fit for each response variable was y ∼ (log10(bio12)*native status) + (bio15*native status). The native status reference level is introduced species, so the estimates for log10(bio12) and bio15 are the slopes for introduced species, and the estimates for the interaction terms are the offset between the native slope and the introduced slope for the two climate variables respectively.

|  |  |  |  |  |
| --- | --- | --- | --- | --- |
| a | Fraction annuals | estimate ± SE | z-statistic | p-value |
|  | (Intercept) | 5.77 ± 0.082 | 70.17 | <0.001 |
|  | log10(bio12) | - 1.98 ± 0.030 | -67.04 | <0.001 |
|  | bio15 | 0.00029 ± 0.0002 | 1.30 | 0.195 |
|  | native status (native) | -2.59 ± 0.08 | -29.31 | <0.001 |
|  | log10(bio12)*native status | 0.12 ± 0.03 | 3.90 | <0.001 |
|  | bio15*native status | 0.013 ± 0.0002 | 53.02 | <0.001 |
| b | Annual richness |  |  |  |
|  | (Intercept) | -1.61 ± 0.10 | -15.68 | <0.001 |
|  | log10(bio12) | 1.45 ± 0.04 | 35.77 | <0.001 |
|  | bio15 | -0.012 ± 0.0003 | -41.63 | <0.001 |
|  | native status (native) | 3.26 ± 0.14 | 23.26 | <0.001 |
|  | log10(bio12)*native status | -1.27 ± 0.06 | -22.79 | <0.001 |
|  | bio15*native status | 0.013 ± 0.0004 | 31.06 | <0.001 |
| c | Perennial richness |  |  |  |
|  | (Intercept) | -3.32 ± 0.11 | -29.79 | <0.001 |
|  | log10(bio12) | 1.96 ± 0.04 | 44.49 | <0.001 |
|  | bio15 | -0.0099 ± 0.0003 | -30.32 | <0.001 |
|  | native status (native) | 3.55 ± 0.15 | 23.38 | <0.001 |
|  | log10(bio12)*native status | -0.97 ± 0.06 | -16.05 | <0.001 |
|  | bio15*native status | 0.0050 ± 0.0005 | 11.07 | <0.001 |

**Supplementary Table 2.** Model fit for effect of population accessibility on likelihood an annual is introduced (versus native). The model fit for each response variable was y ∼ (log10(bio12)*population accessibility) + (bio15* population accessibility). WorldClim climate variable bio12 is mean annual precipitation (mm) and bio15 is precipitation seasonality.

| Parameter | estimate ± SE | z-statistic | p-value |
| --- | --- | --- | --- |
| (Intercept) | -7.30 ± 0.10 | -74.19 | <0.001 |
| population accessibility | 5.33 ± 0.13 | 42.32 | <0.001 |
| log10(bio12) | 2.57 ± 0.04 | 61.95 | <0.001 |
| bio15 | -0.03 ± 0.000342 | -74.42 | <0.001 |
| pop_accessibility:log10(bio12) | -1.89 ± 0.05 | -41.24 | <0.001 |
| pop_accessibility:bio15 | 0.03 ± 0.000539 | 46.58 | <0.001 |

**Supplementary Table 3.** Relationships between climate variables and species richness and fractional richness, using resampled data from the rarefaction analysis. Responses of a. fraction annuals; b. annual richness; and c. perennial richness to log10(bio12) (mean annual precipitation, mm), bio15 (annual precipitation seasonality), and native status (native, introduced). The model fit for each response variable was y ∼ (log10(bio12)*native status) + (bio15*native status). The native status reference level is introduced species, so the estimates for log10(bio12) and bio15 are the slopes for introduced species and the estimates for the interaction terms are the offset between the native slope and the introduced slope for the two climate variables respectively. WorldClim climate variable bio12 is mean annual precipitation (mm) and bio15 is precipitation seasonality.

|  |  |  |  |  |
| --- | --- | --- | --- | --- |
| a | Fraction annuals | estimate ± SE | z-statistic | p-value |
|  | (Intercept) | 4.69 ± 0.19 | 24.40 | <0.001 |
|  | log10(bio12) | -1.45 ± 0.07 | -20.09 | <0.001 |
|  | bio15 | -0.0065 ± 0.0005 | -13.18 | <0.001 |
|  | native status (native) | -1.67 ± 0.20 | -8.29 | <0.001 |
|  | log10(bio12)*native status | -0.35 ± 0.076 | -4.59 | <0.001 |
|  | bio15*native status | 0.019 ± 0.0005 | 35.34 | <0.001 |
| b | Annual richness |  |  |  |
|  | (Intercept) | -0.19 ± 0.088 | -2.13 | 0.034 |
|  | log10(bio12) | 0.59 ± 0.035 | 16.85 | <0.001 |
|  | bio15 | -0.007 ± 0.0003 | -27.78 | <0.001 |
|  | native status (native) | 3.71 ± 0.12 | 30.83 | <0.001 |
|  | log10(bio12)*native status | -1.43 ± 0.05 | -29.89 | <0.001 |
|  | bio15*native status | 0.014 ± 0.0004 | 37.44 | <0.001 |
| c | Perennial richness |  |  |  |
|  | (Intercept) | -1.69 ± 0.08 | -20.42 | <0.001 |
|  | log10(bio12) | 0.99 ± 0.03 | 30.35 | <0.001 |
|  | bio15 | -0.003 ± 0.0002 | -12.55 | <0.001 |
|  | native status (native) | 3.69 ± 0.11 | 32.69 | <0.001 |
|  | log10(bio12)*native status | -0.96 ± 0.04 | -21.43 | <0.001 |
|  | bio15*native status | 0.004 ± 0.0003 | 11.30 | <0.001 |

**Supplementary Table 4.**
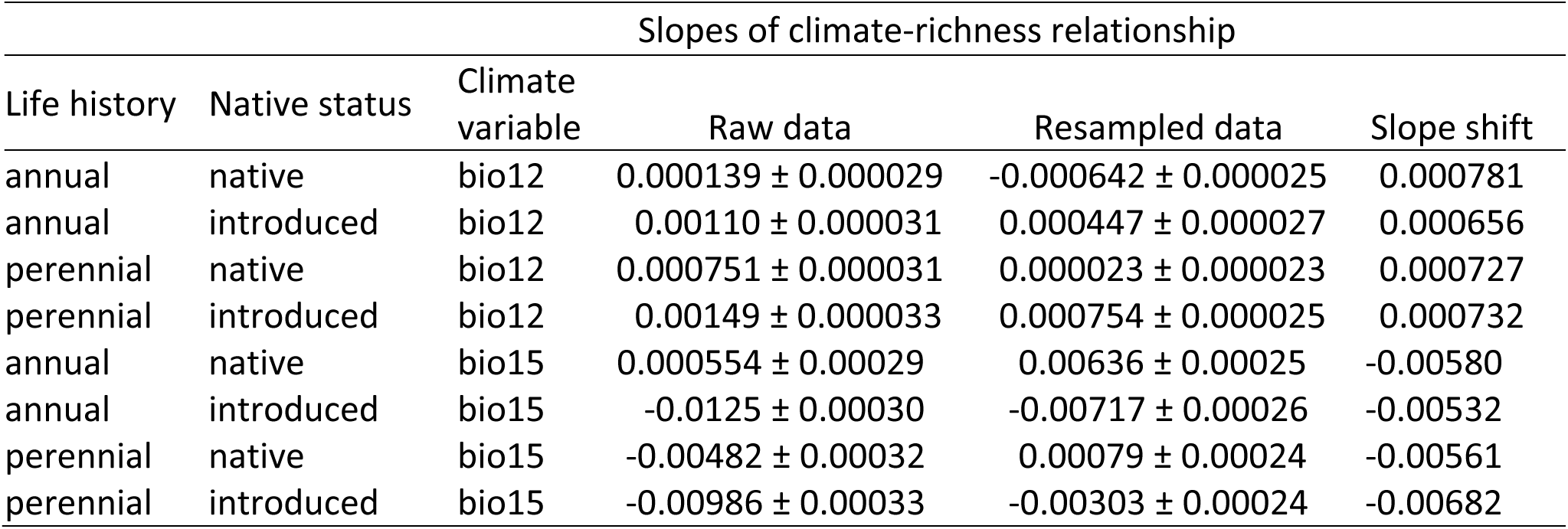
Comparison of slopes for raw (Table S1) and resampled data from the rarefaction analysis (Table S3), including the shift in slopes between the two datasets. WorldClim climate variable bio12 is mean annual precipitation (mm) and bio15 is precipitation seasonality. The model fit for each life history group was richness ∼ (log10(bio12)*native status) + (bio15*native status).

**Supplementary Fig. 1.**
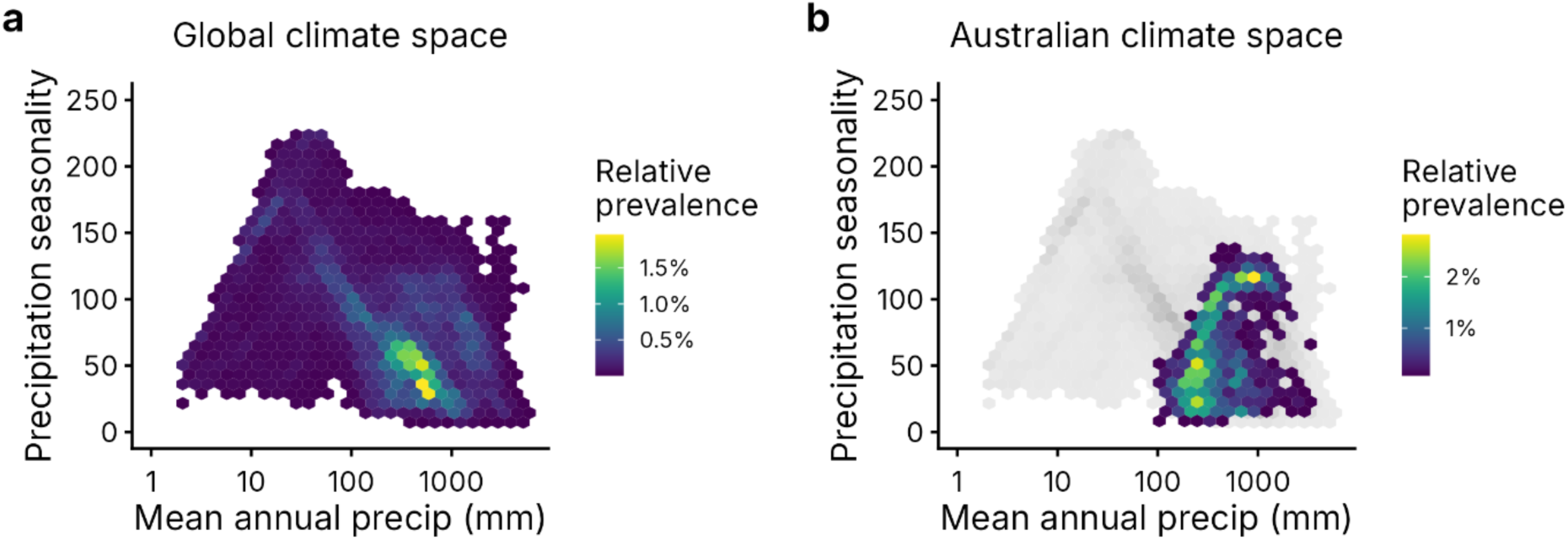
a. Global climate space and b. Australia’s climate space in comparison to global climate space.

**Supplementary Fig. 2.**
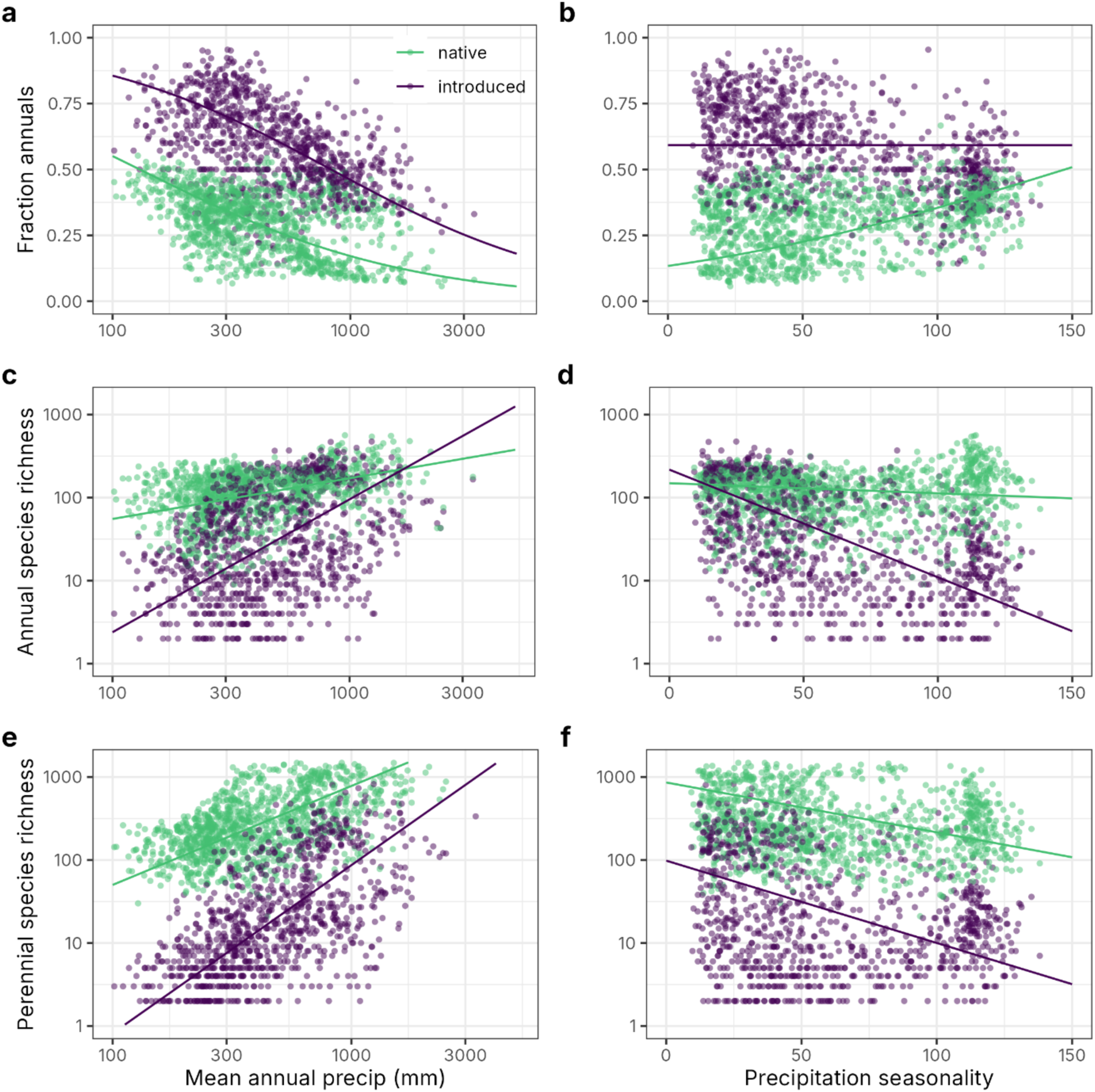
Figure 3 repeated, with grid cells with fewer than 100 records omitted.

**Supplementary Fig. 3.**
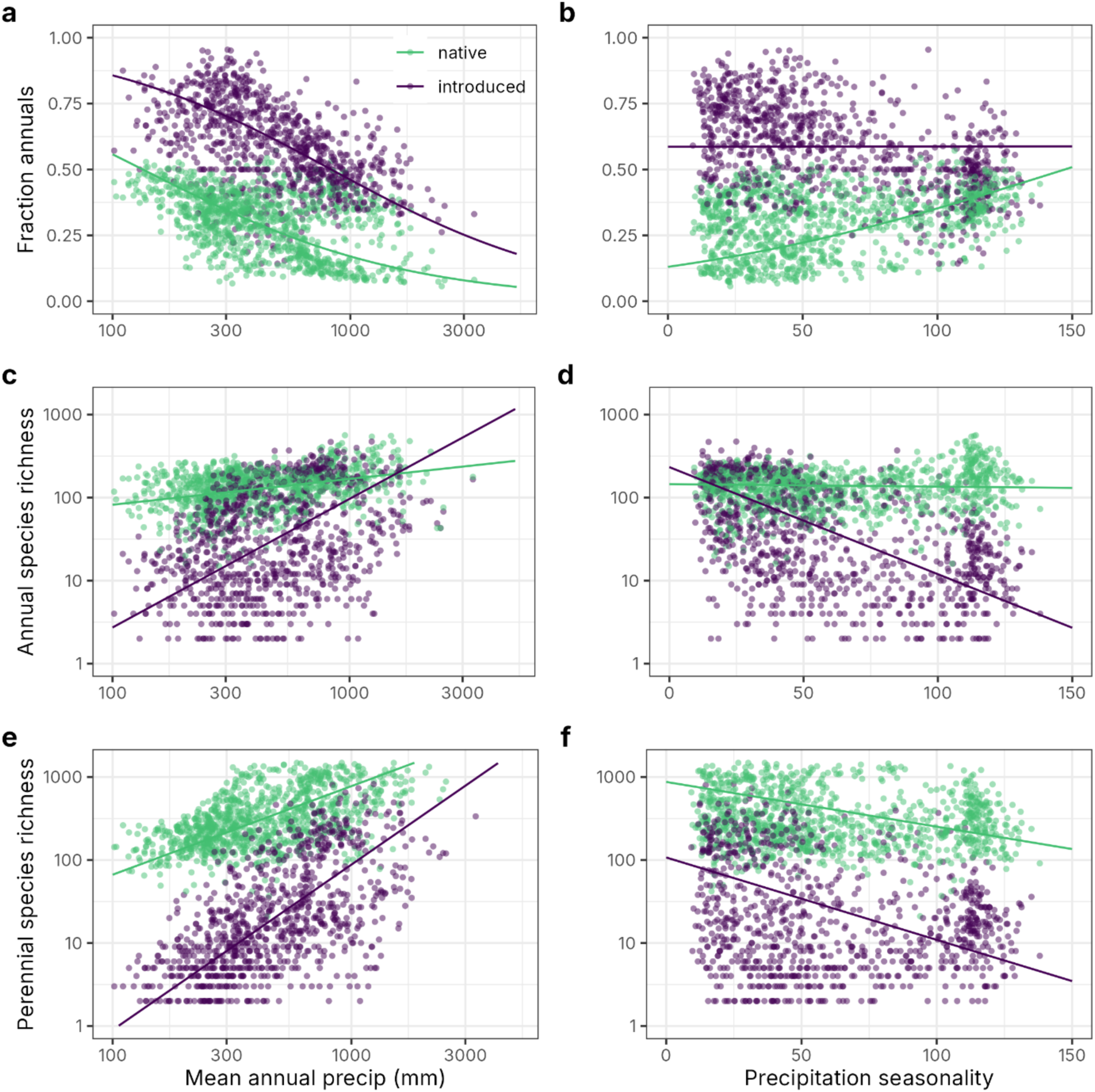
Figure 3 repeated, with grid cells with fewer than 250 records omitted.

**Supplementary Fig. 4.**
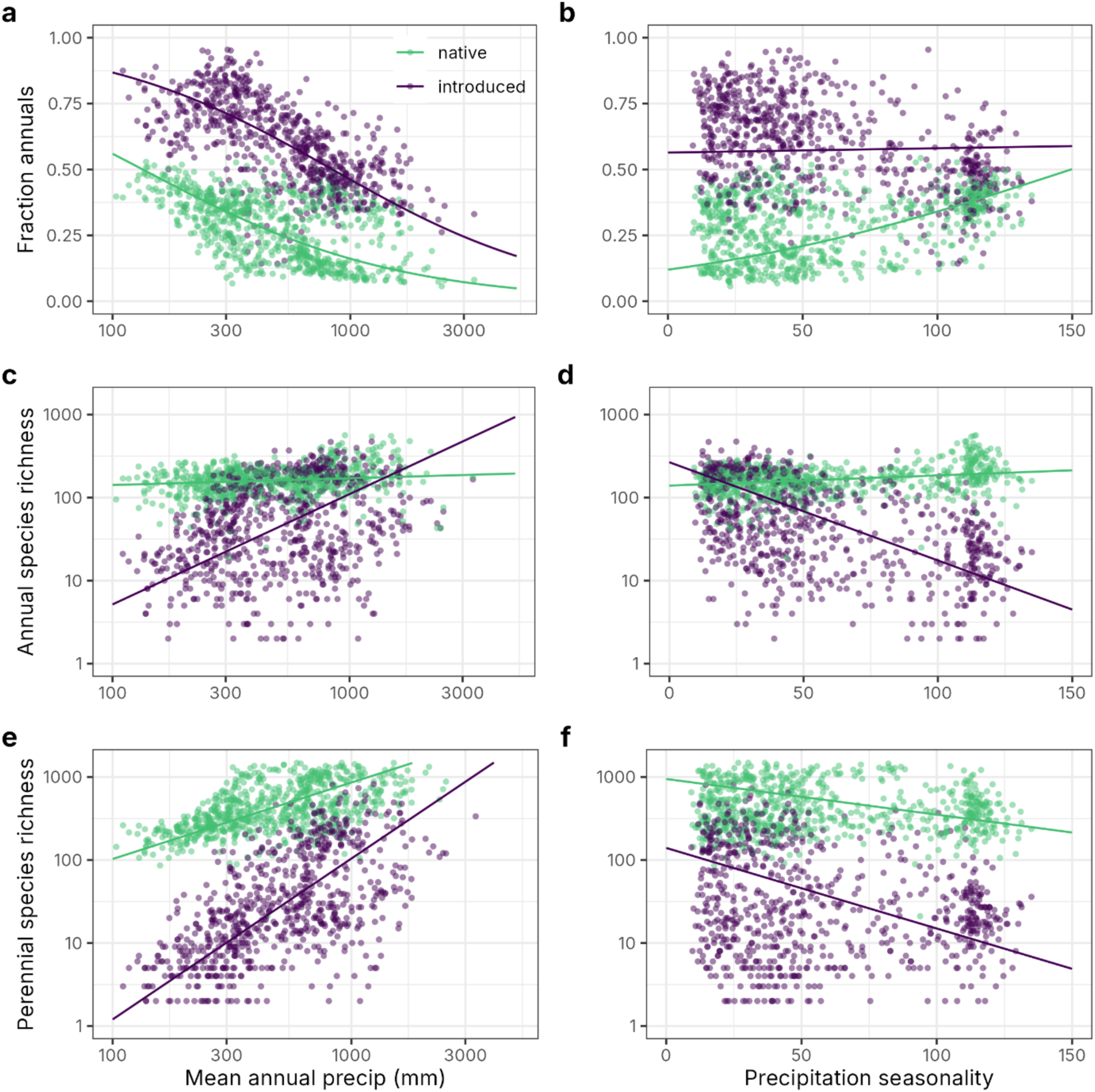
Figure 3 repeated, with grid cells with fewer than 1000 records omitted.

**Supplementary Fig. 5.**
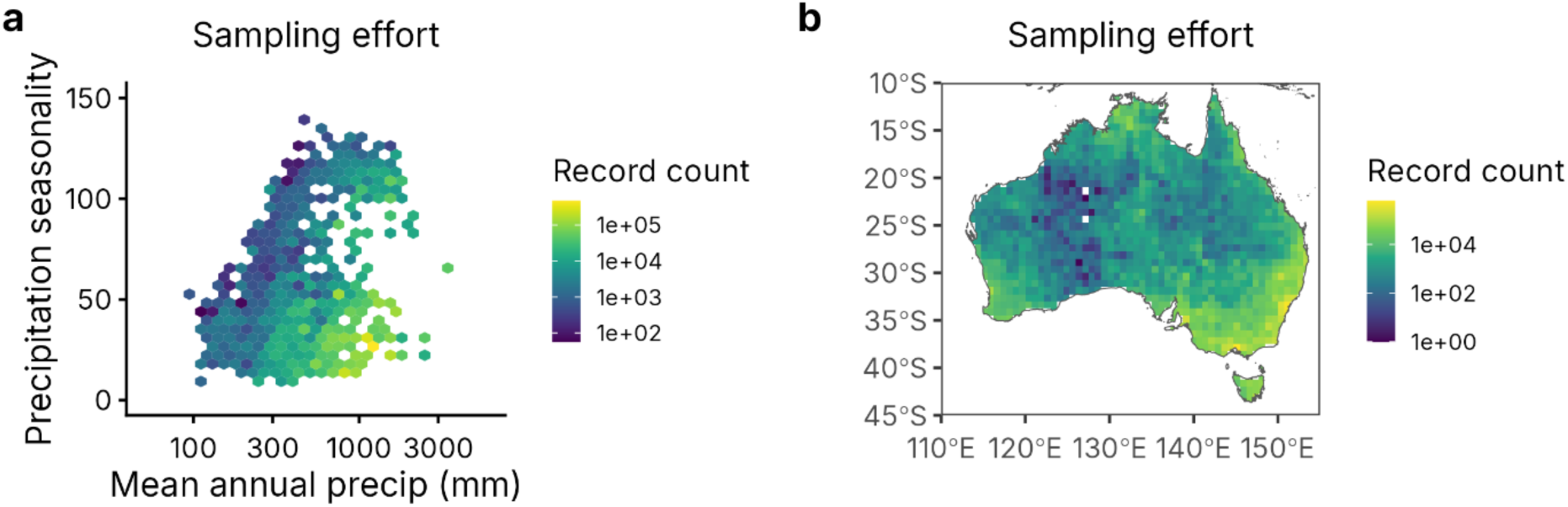
Shifts in sampling effort across a. Australian climate space; and b. the Australian land surface.

**Supplementary Fig. 6.**
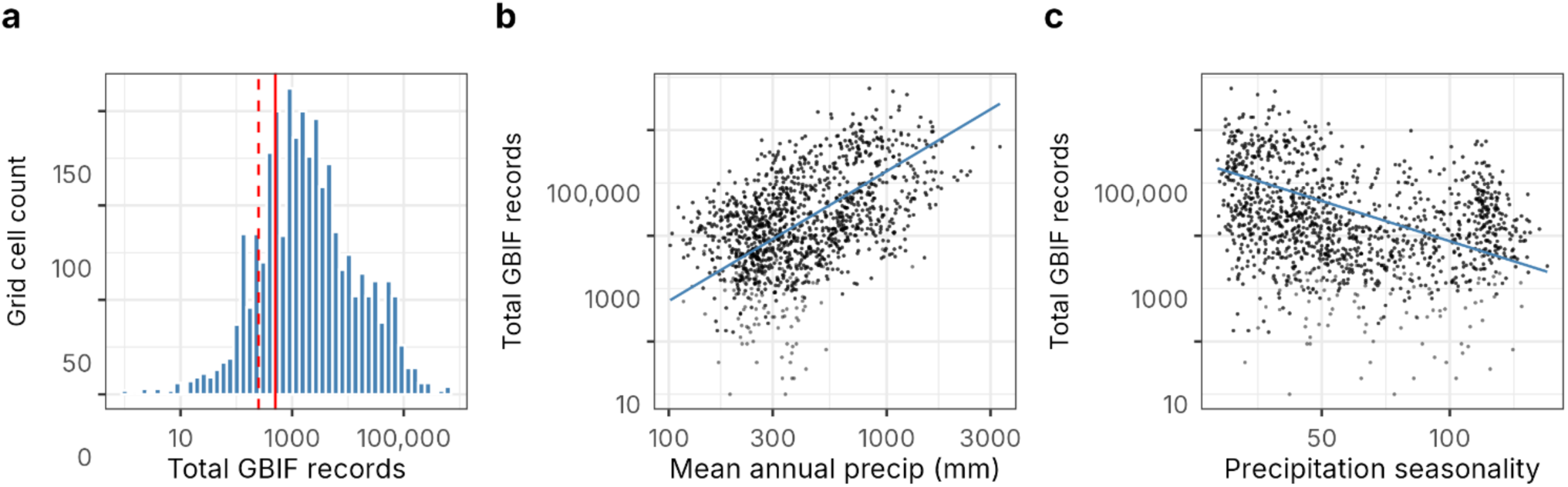
Shifts in sampling effort across climate gradients, including a. sampling effort distribution, b. sampling effort versus mean annual precipitation (mm), and c. sampling effort versus precipitation seasonality.

